# Semidilute Polymer Scaling Controls PEG-induced Protein Phase Separation

**DOI:** 10.64898/2026.10.25.761614

**Authors:** Mohammad Ahmad, Hala Ouzon-Shubeita, Malvern Whende, Alessandro Ianiro, Aleksander A. Rebane

## Abstract

When does polyethylene glycol (PEG) mass concentration provide a common measure of crowding strength across polymer chain lengths? Here we use bovine serum albumin (BSA) and PEG as a controlled depletion system and measure both coexisting phases to determine binodals, tie lines, and critical points across PEG molecular weights of 4–100 kDa. Phase diagrams for PEG 20, 35, and 100 kDa approximately collapse in mass-concentration coordinates, consistent with semidilute polymer scaling. Generalized free volume theory (GFVT) describes the BSA binodals and tie lines. It also approximately describes binodals for three additional globular proteins in PEG 35K, extending the comparison across proteins spanning 7.2–66.4 kDa. For the binary PEG mixtures tested, GFVT approximates the binodals using an effective polymer size corresponding to the weight-average molecular weight. These results identify a regime in which PEG mass concentration (g/L) provides a transferable measure of depletion-induced crowding.

## INTRODUCTION

Biomolecular phase separation driven by weak multivalent interactions is a central mechanism for organizing intracellular material into condensates such as P granules, nucleoli, stress granules, and synaptic densities [1–7]. Because the interactions that stabilize condensates are often weak and collective, condensate formation, dissolution, composition, and material properties are sensitive to environmental variables such as salt, osmolytes, temperature, pH, and macromolecular crowding [8–12]. Crowding is especially important in the cytoplasm, where up to 30% volume fraction is occupied by macromolecules [13]. This restricts the free volume available for biomacromolecules and gives rise to entropic depletion attractions between proteins, nucleic acids, and membranes [14–17].

Depletion occurs when inert macromolecules (depletants) are excluded from the vicinity of a larger solute [18]. The region of exclusion surrounding the solute is called the depletion layer. When two solutes approach close enough such that their depletion layers overlap, the volume accessible to the depletant increases. This, in turn, increases depletant entropy and produces an effective attraction between the larger solutes [19, 20]. Depletion can therefore promote biomolecular association, membrane adhesion, and segregative liquid-liquid phase separation (LLPS) even in the absence of direct attractive interactions [21–27].

In reconstituted phase separation assays, macromolecular crowding is often mimicked by adding nominally inert polymers such as dextran, Ficoll, or polyethylene glycol (PEG) [28]. PEG is particularly widely used because it can strongly shift protein phase boundaries in a concentration- and molecular-weight-dependent manner [29, 30]. However, PEG concentration alone is not generally a transferable measure of crowding strength: changing PEG molecular weight changes both the size of an isolated polymer coil and the concentration at which polymer coils begin to overlap. As a result, the same mass concentration of PEG can represent distinct physical regimes for different molecular weights. This ambiguity complicates comparisons across condensate assays and motivates a quantitative criterion for when PEG mass concentration can, and cannot, be used as a common crowding variable. Here, we use BSA-PEG phase separation as a controlled model system to identify this criterion. BSA is a compact globular protein, and at pH 7 it carries a net negative charge (pI ≈4.8 ±0.1), while PEG is a highly hydrated neutral polymer. Under the conditions used here, BSA and PEG undergo segregative phase separation into a BSA-rich condensate phase and a PEG-rich supernatant phase (Fig. 1A), providing a controlled system for testing a depletion-based description of protein phase separation. A practical advantage of this system is that BSA and PEG can be used in bulk quantities, which permits direct measurement of full phase diagrams, including binodals, tie lines, and critical points (Fig. 1B).

**FIG. 1.**
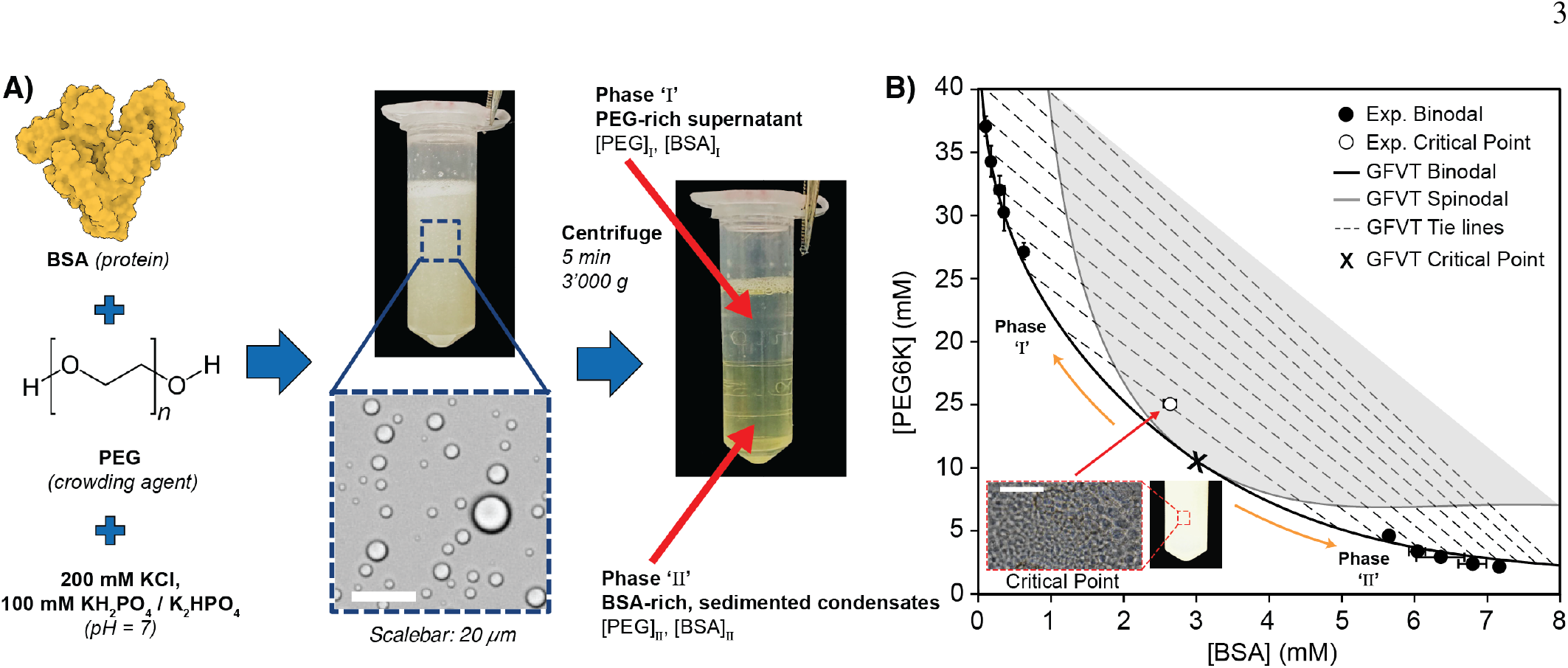
Measurements of both coexisting phases test GFVT descriptions of BSA-PEG binodals and tie lines. **A)** Schematic of liquid-liquid phase separation of BSA driven by PEG-induced depletion interactions. The binodal and tie lines are measured by separating the BSA-rich condensate phase from the PEG-rich supernatant phase and quantifying their compositions using UV-VIS spectroscopy and proton NMR. **B)** Experimentally measured binodal (*black circles*) and critical point (*open circle*) for BSA in the presence of PEG 6K. GFVT, with *a* = 3.0 nm and *R*_*g*_ = 2.45 nm (*q* = 0.82), predicts binodals (*black solid*), tie-lines (*gray dashed*), spinodal (*gray solid*), and critical point (*cross*). Inset shows the critical point. Scale bar, 20 µm. BSA structure PDB ID: 1E7I [31].

We combine measurements of both coexisting phases with generalized free volume theory (GFVT) to identify when PEG mass concentration provides a common crowding variable across polymer chain lengths. These measurements determine both arms of the binodal and the tie lines connecting them, directly constraining protein and PEG partitioning. In the dilute-chain regime, the relevant depletion length is set by the radius of gyration of individual PEG coils, so phase boundaries depend strongly on PEG molecular weight. In the semidilute regime, overlapping PEG coils form a polymer mesh, and the correlation length of this mesh becomes the relevant depletion length. Consistent with this scaling, the BSA-PEG phase diagrams for PEG 20K, 35K, and 100K approximately collapse when plotted against PEG mass concentration. For the defined binary PEG mixtures tested, BSA binodals are approximated by those of monodisperse PEG with the same weight-average molecular weight. We also examine whether GFVT describes ovalbumin, maltose-binding protein (MBP), and GB1 binodals in PEG 35K using literature-based protein radii and the same PEG radius of gyration calibrated against BSA. Where the literature reports a range of protein radii, we select a value within that range based on agreement with the binodal. Together, these results identify conditions under which PEG mass concentration (g/L) provides a transferable measure of crowding strength across PEG molecular weights and support the applicability of a common depletion-based description across globular proteins.

## RESULTS

### Direct coexistence measurements test GFVT for BSA-PEG

Figure 1A summarizes the experimental approach used to determine BSA-PEG phase diagrams. Mixtures of BSA and PEG 6K were prepared in buffered solution (100 mM potassium phosphate at pH 7, 200 mM KCl), where they underwent LLPS into a dense BSA-rich condensate phase and a dilute PEG-rich supernatant phase [32]. After centrifugation, macroscopic quantities of the two coexisting phases were collected and their compositions were measured directly: BSA concentration was determined by UV absorbance, and PEG concentration was determined by ^1^H NMR. Repeating this procedure over a range of starting compositions yielded both arms of the binodal and the tie lines connecting the compositions of each coexisting pair. These measurements directly quantify PEG partitioning between the phases, providing constraints beyond the onset of phase separation. We determined the critical point by the method of equal volumes, in which phase-separated samples were iteratively diluted at approximately equal phase volumes until the density difference between the two phases became too small to permit further separation by centrifugation [12, 33, 34].

To interpret these phase diagrams, we used GFVT [35]. In the model, BSA is represented as a colloidal hard sphere of radius *a*, and PEG is represented as a polymer depletant with radius of gyration *R*_*g*_. The relevant size ratio is

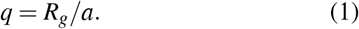

GFVT yields the Helmholtz free energy, osmotic pressure, and protein chemical potential as functions of the BSA volume fraction, *η* = 4*πa*^3^*n*_*η*_ */*3, and the PEG volume fraction, 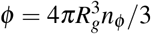, where *n*_*η*_ and *n*_*φ*_ are the number densities of BSA and PEG, respectively. For a given *q*, the model predicts the liquid-liquid coexistence binodal, tie lines, spinodal, and critical point (see Materials and Methods).

We fixed *a* = 3.0 nm, corresponding to the crystallographic radius of BSA, and fitted *R*_*g*_ to the measured binodal. For PEG 6K, the best-fit value *R*_*g*_ = 2.45 nm gave quantitative agreement with the measured binodal and tie lines (Fig. 1B). This value is consistent with the expected radius of gyration of PEG 6K in water. The GFVT critical point occurred at lower PEG concentration and higher BSA concentration than the measured critical point. GFVT is a mean-field theory and does not capture the long-wavelength concentration fluctuations near critical points in the 3D Ising universality class [36]. The critical-point discrepancy may reflect this limitation or the the hard-sphere representation of BSA. Away from the immediate critical region, agreement with the binodal and tie lines supports a depletion-based description of BSA-PEG coexistence under the conditions studied.

### PEG molecular weight shifts phase boundaries

We next measured BSA phase diagrams in the presence of PEG 4K, 6K, 8K, 20K, 35K, and 100K (Fig. 2A), with representative condensate images shown in Fig. 2C. Increasing PEG molecular weight shifted the binodal to lower molar concentrations of both BSA and PEG. The measured critical point likewise moved to lower BSA and PEG concentrations as PEG molecular weight increased. This behavior is consistent with the notion that larger polymer coils lead to larger depletion layers around each BSA molecule and therefore generate stronger effective attractions at lower molar concentration. We captured this trend with GFVT by changing only the polymer size parameter *R*_*g*_, while keeping the BSA radius fixed, yielding the critical line (Fig. 2A, *gray dashed*), which shows how the critical point shifts with *q*.

**FIG. 2.**
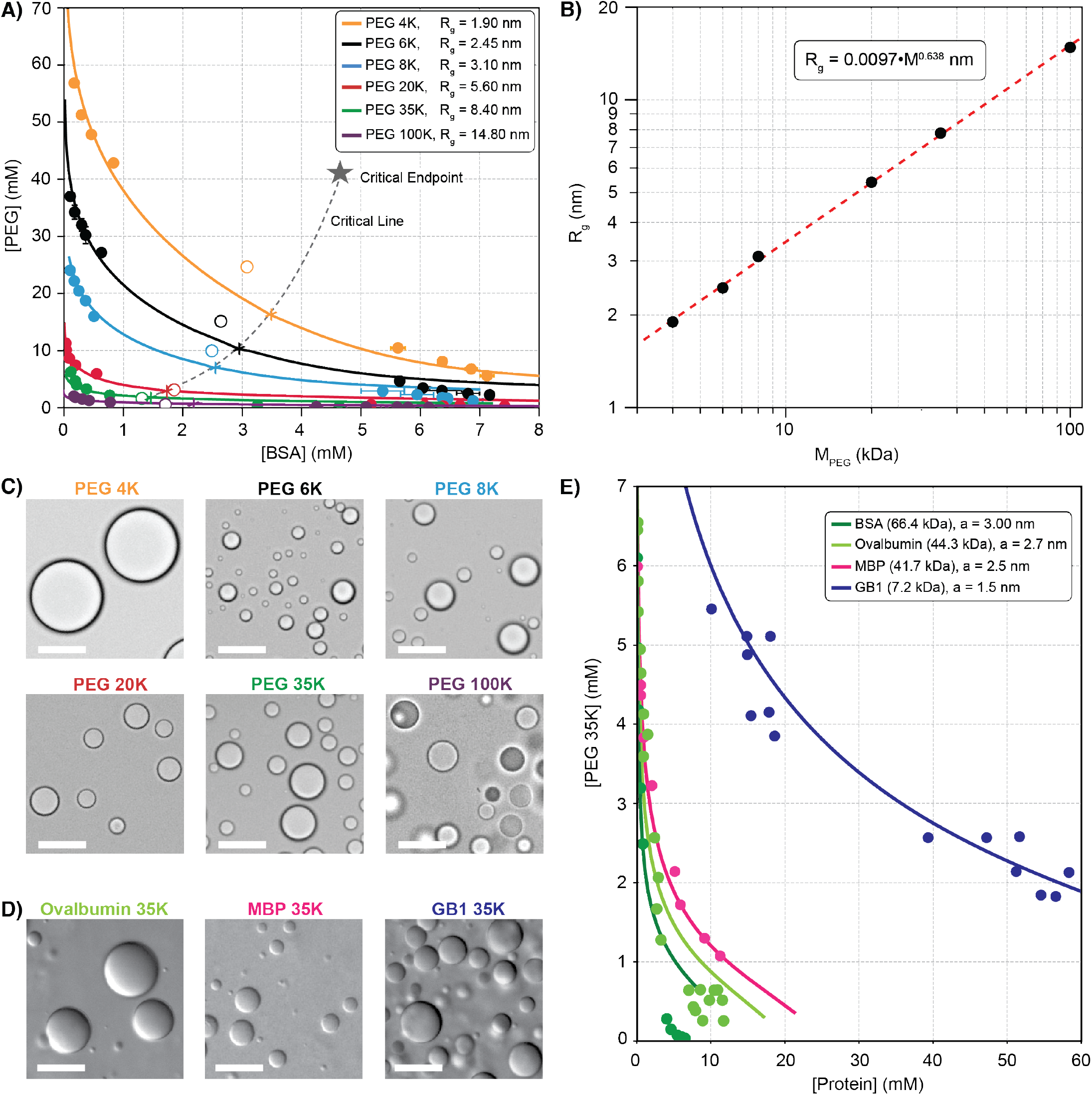
GFVT describes protein-PEG phase diagrams across polymer and protein sizes. **A)** Experimentally measured binodals (*circles*) and critical points (*open circles*), and GFVT binodals (*curves*) and critical points (*crosses*) for BSA in the presence of PEG 4K (*q* = 0.63, *yellow*), PEG 6K (*q* = 0.82, *black*), PEG 8K (*q* = 1.03, *blue*), PEG 20K (*q* = 1.87, *red*), PEG 35K (*q* = 2.80, *green*), and PEG 100K (*q* = 4.93, *purple*). The critical line (*gray, dashed*) shows how the critical point moves to higher concentrations with decreasing PEG molecular weights up to the critical endpoint (*star*) where liquid-liquid coexistence becomes metastable. **B)** Log-log plot of the radius of gyration of PEG versus the molecular weight of PEG in the GFVT binodals shown in A). Brightfield microscopy images of **C)** BSA condensates in the presence of varying sizes of PEG and of **D)** ovalbumin, maltose-binding protein (MBP), and GB1 condensates in the presence of PEG 35 K. Scale bar, 20 µm. **E)** Experimentally measured binodals (*symbols*) and GFVT binodals (*solid curves*) for BSA (*green*), ovalbumin (*light green*), MBP (*pink*), and GB1 (*dark blue*), each in the presence of PEG 35K. All GFVT curves in E) use the same PEG radius of gyration as the BSA-PEG 35K calculation in A).

It is intuitive that macromolecules can induce crowding whereas small molecules such as ethylene glycol or glucose cannot. What is the size cutoff beyond which a macro-molecule becomes too small to act as a crowder? GFVT predicts a lower size ratio below which liquid-liquid coexistence becomes metastable with respect to a solid phase. This so-called critical endpoint occurs at approximately *q* = 0.388. Below this value, the depletion attraction is too short-ranged to stabilize liquid-liquid coexistence against crystallization or aggregation into a solid-like precipitate [37]. For the BSA system studied here, this endpoint corresponds to PEG with molecular weight of approximately 1.8 kDa (Fig. 2A, *star*). Consistent with this prediction and with previous experiments [38], PEG molecular weights below 2 kDa did not produce stable BSA LLPS at room temperature.

The fitted PEG radii of gyration followed a power-law relationship with molecular weight,

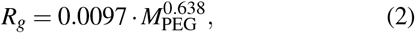

where *R*_*g*_ is in nm and *M*_PEG_ is in Da (Fig. 2B). The empirical Flory exponent *ν* = 0.638 lies within the reported range of 0.58-0.67 for PEG in water [39, 40]. This fitted exponent describes the molecular-weight dependence of the inferred PEG radii and is distinguished below from the asymptotic good-solvent exponent used in semidilute scaling. The consistency with reported coil-size scaling supports the interpretation that polymer size contributes strongly to the observed shifts in phase boundaries. More detailed models could include BSA charge and solvent electrostatics in addition to polymer excluded-volume interactions [41–44], but the present GFVT comparison uses one fitted PEG radius of gyration for each PEG molecular weight, with the BSA radius held fixed.

To test whether the GFVT description extends beyond BSA, we measured binodals for three additional globular proteins in the presence of PEG 35K, including ovalbumin (44.3 kDa), maltose-binding protein (MBP; 41.7 kDa), and B1 domain of streptococcal protein G (GB1; 7.2 kDa). All three proteins form spherical condensates in the presence of sufficient PEG 35K (Fig. 2D), and their experimental binodals approximately follow the GFVT predictions (Fig. 2E). The calculations used protein radii *a* = 2.7 nm for ovalbumin [45], *a* = 2.5 nm for MBP [46], and *a* = 1.5 nm for GB1 [47], which were independently determined based on the literature and if a range was given, the value was chosen within that range that best fitted the binodal. The PEG radius of gyration was held at the value used for BSA-PEG 35K, without refitting for the other proteins. Binodals for PEG 35K plotted with expanded PEG-concentration-axis compared to Fig. 2A reveal that GFVT systematically overestimates the dense phase PEG concentration for BSA and ovalbumin, but not MBP or GB1. This could be due to BSA’s and ovalbumin’s propensity to oligomerize [48]. Nevertheless, approximate agreement with the measured binodals therefore supports the transferability of the GFVT description across proteins spanning 7.2–66.4 kDa, without adjusting either size parameter to the additional binodal data.

### A semidilute scaling transition collapses the phase diagrams

Plotting the phase diagrams in units of *g/L* reveals two regimes of PEG-induced crowding (Fig. 3A). At PEG molecular weights below 20 kDa, the phase diagram is sensitive to molecular weight. This is the dilute-chain regime, where PEG is below the polymer overlap concentration and individual PEG coils act as distinct depletants. The depletion length is set by the coil size, *δ* ~*R*_*g*_, whose molecular-weight dependence is given by Eq. 2. In contrast, the phase diagrams for PEG molecular weights of 20 kDa and above collapse on a single curve. This behavior is consistent with coexistence in the semidilute regime, where PEG is above the polymer overlap concentration and the depletion length is set by the correlation length of the polymer mesh, *δ*~ *ξ*, rather than the the size of an isolated chain. For a given protein, this scaling predicts that phase boundaries become less sensitive to PEG chain length once the relevant coexistence region lies in the semidilute regime.

**FIG. 3.**
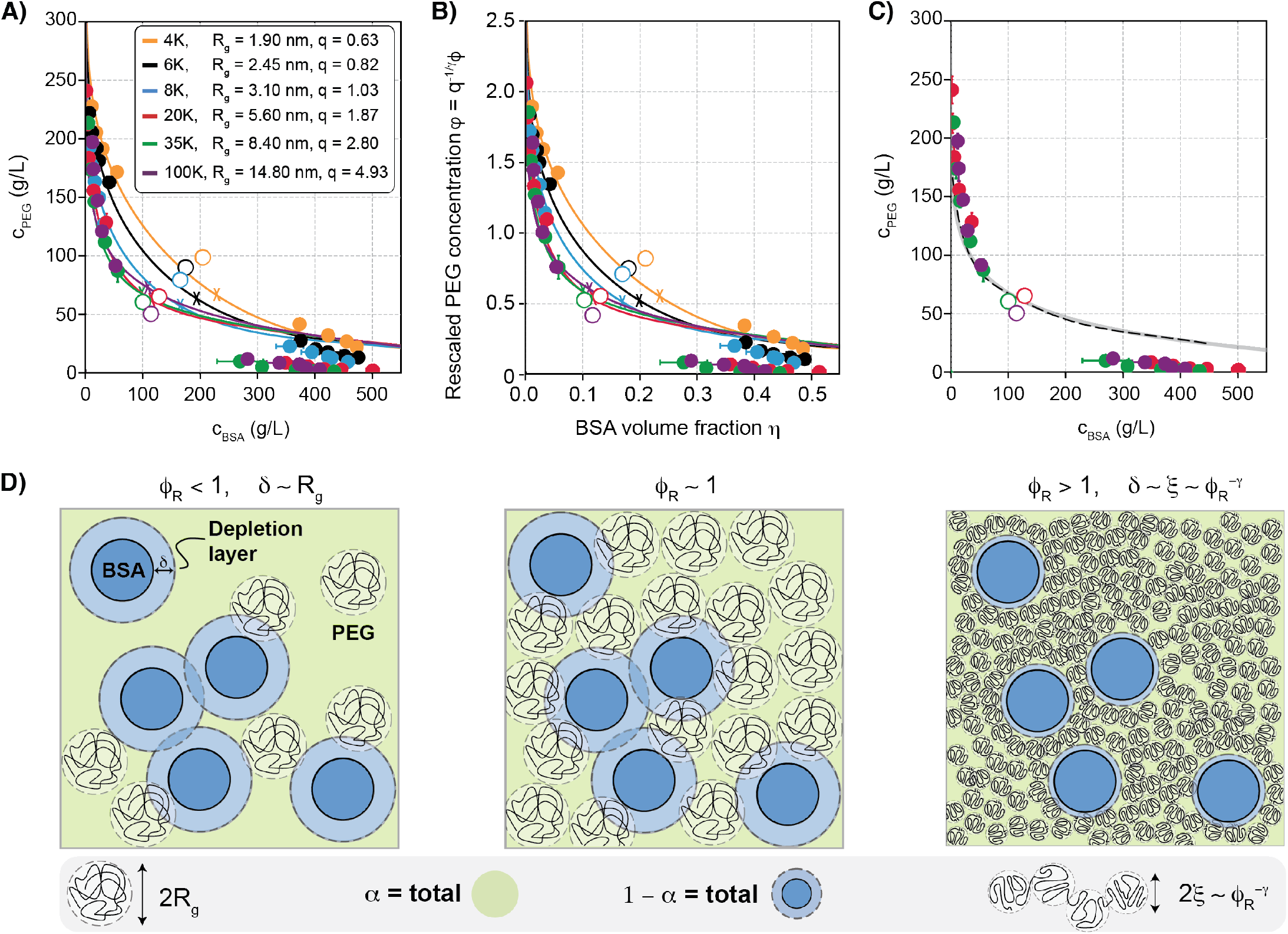
Semidilute scaling transition collapses BSA-PEG phase diagrams. **A)** BSA-PEG phase diagrams plotted in concentration units of grams per liter. Shown are experimentally measured binodals (*circles*) and critical points (*open circles*) and GFVT predictions of binodals (*solid curves*) and critical points (*crosses*). **B)** Binodals and critical points for different PEG molecular weights plotted with respect to the rescaled PEG concentration, *ϕ* = *q*^−1*/γ*^ *φ*, and BSA volume fraction, *η*. **C)** Analytical approximation of binodal (*dashed*) and exact solution (*gray*) for *q* = 16 overlain with experimentally measured binodal for PEG 20K, PEG 35K, and PEG 100K. **D)** Schematic illustrating the definition of *φ*_*R*_ as the fraction of free volume, ⟨*V*_free_⟩ = *α*· *V* (total area shaded in green), occupied by PEG. The total excluded volume due to BSA and its depletion layer (area shaded in blue) is ⟨*V*_ex_⟩ = (1− *α*) ·*V*. Phase diagrams are shown for PEG 4K (*yellow*), PEG 6K (*black*), PEG 8K (*blue*), PEG 20K (*red*), PEG 35K (*green*), and PEG 100K (*purple*).

GFVT makes this expectation quantitative. The PEG concentration relevant to chain overlap is its concentration within the accessible free volume. The free volume fraction *α* (Fig. 3D, *green*) is the fraction of total volume occupied by neither BSA (*dark blue*) nor its surrounding depletion layer (*light blue*). The relative PEG concentration in this volume is

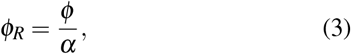

with chain overlap occurring at approximately *φ*_*R*_ = 1. In the semidilute regime (*φ*_*R*_ *>* 1), PEG chains overlap, resulting in an approximately ideal solution of correlation blobs whose size is given by the correlation length,

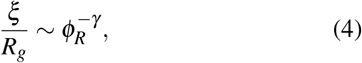

with *γ* = 0.77 the De Gennes exponent for a polymer in good solvent [49, 50]. The polymer osmotic pressure scales with the number density of correlation blobs, giving

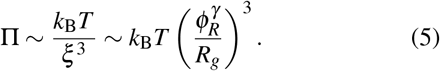

Since *q* = *R*_*g*_*/a*, the corresponding dimensionless pressure scales as [51]

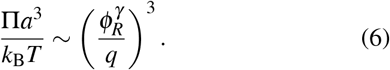

Consequently, in the large-*q* limit, rescaling the PEG concentration using

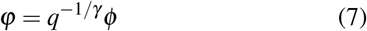

gives a common asymptotic form for the BSA-PEG binodals (Fig. 3B). This scaling has been predicted for polymer-colloid mixtures [52] and supported by simulations [53] and experiments with microemulsion-polymer mixtures [54, 55].

The relation between the rescaled concentration and PEG mass concentration depends on the Flory exponent. At fixed protein radius, *R*_*g*_ ~ *M*^*ν*^ and 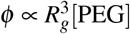 give

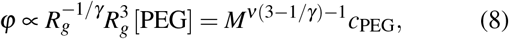

where *c*_PEG_ = *M*[PEG]. In the semidilute regime, the relation

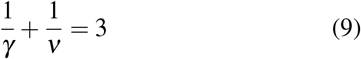

makes *ϕ* proportional to *c*_PEG_. The value *γ* = 0.77 corresponds to *ν*≈ 0.588 under this relation. Combining *γ* = 0.77 with the empirical exponent *ν*_fit_ = 0.638 instead leaves a residual factor *M*^0.085^, so the equivalence is approximate for the parameterization used here. Experimentally, BSA-PEG phase diagrams for PEG 20K, 35K, and 100K approximately overlap when plotted in g/L, whereas lower-molecular-weight PEGs retain molecular-weight-dependent phase boundaries. The binodals for higher molecular weights can be approximated by a common analytical curve (Fig. 3C, *dashed*; see Materials and Methods).

In polymer solution notation, the same collapse can be described in terms of polymer volume fraction, *f* = *Nν*_*s*_*ρ*, where *N* is the number of monomers per PEG chain, *ν*_*s*_ is the volume per monomer, and *ρ* is the polymer chain number density. Since *N* is proportional to the PEG molecular weight, it follows that *f* ∝ *ϕ* and the binodals for large molecular weights collapse when plotted in terms of *f*. The collapse therefore identifies the physical condition under which PEG mass concentration or polymer volume fraction can serve as effective crowding coordinates.

### Thermodynamic origin of the semidilute collapse

The scaling collapse can be understood by examining the depletion length, osmotic pressure, free volume fraction, and BSA chemical potential along the GFVT binodals. The measured PEG concentration in the full sample volume (Figs. 1B and 2A) is related to the relative polymer concentration in the accessible free volume by

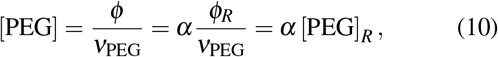

where 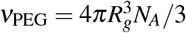 and *N*_*A*_ is Avogadro’s constant.

The free volume fraction depends on BSA concentration and the thickness of the depletion layer, *δ* (Fig. 3D). For a polymer in good solvent, we approximate the curvature-corrected depletion thickness around a spherical protein by interpolating between the dilute-chain- and semidilute-limits according to [51, 56, 57]

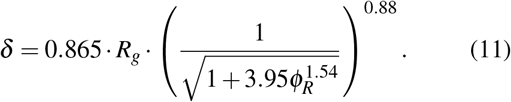

Figure 4A shows how *δ* decreases with increasing PEG concentration (*solid curves*). Thick curves indicate the regions corresponding to the binodals in Fig. 2A. For [PEG]_*R*_ ≥20 mM, the depletion length decreases with increasing PEG molecular weight. Along the BSA-PEG binodals, *δ* ranges from 0.5 nm to 1.5 nm (*thick curves*), regardless of molecular weight, with the greatest depletion length at the critical point (*circles*). The depletion length asymptotically approaches the polymer correlation length for PEG above the overlap concentration (*dashed*). In fact, for PEG with molecular weights greater than 20 kDa, binodals lie entirely in the semidilute regime and 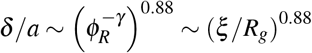.

**FIG. 4.**
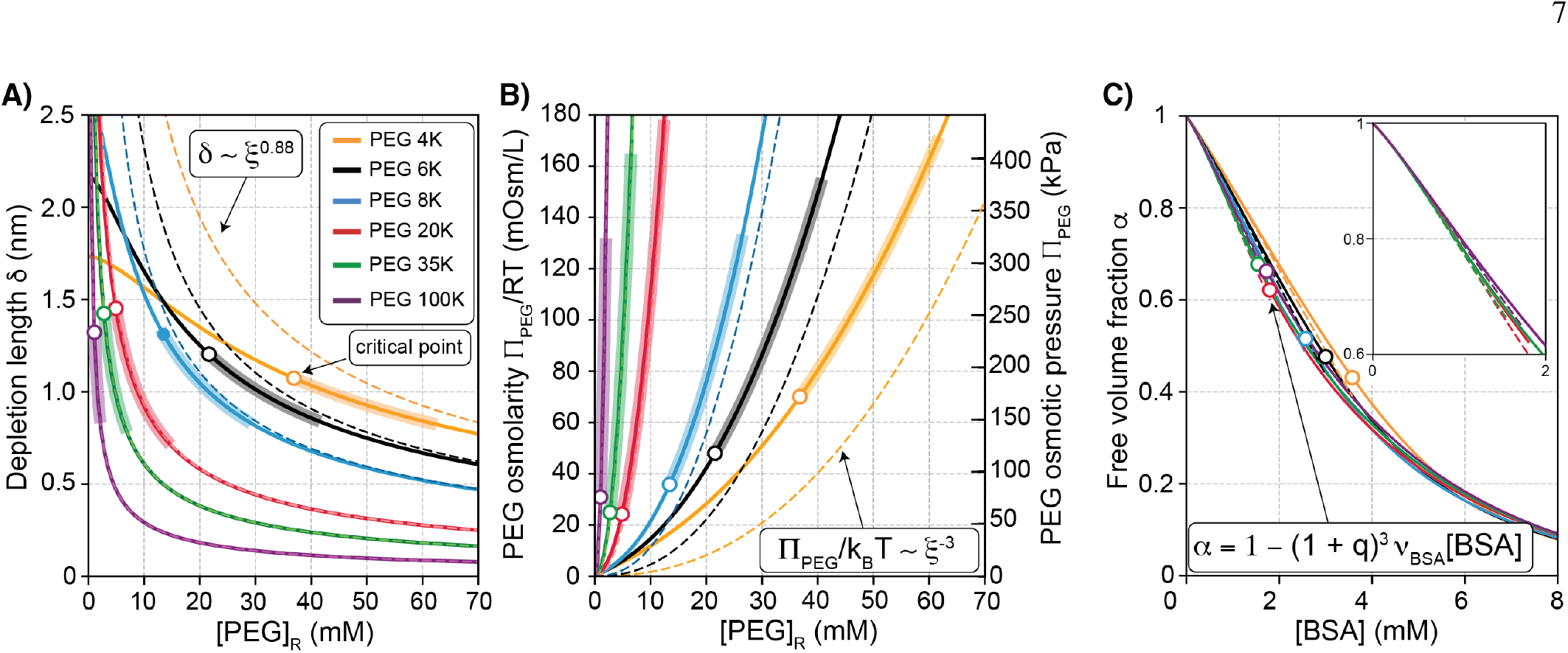
Depletion length, osmotic pressure, and free volume along BSA-PEG binodals. **A)** Depletion length, *δ* as a function of PEG concentration (*solid curves*), with its semidilute approximation *δ*~ *ξ* ^−0.88^ (*dashed*). Two-phase coexistence regions are marked by thick curves. The molar PEG concentration is related to the volume fraction of PEG via [PEG]_*R*_ = *φ*_*R*_*/ν*_PEG_, where *ν*_PEG_ is the molar volume of PEG. **B)** Contribution of PEG to osmotic pressure (*solid curves*) and the semidilute approximation Π~*k*_B_*T/ξ* ^3^ (*dashed*). **C)** Free volume fraction, *α*, as a function of BSA concentration along the binodal (*solid curves*). The supernatant arm of the binodal lies to the left of the critical point (*circle*), and the condensate arm lies to its right. For low BSA concentrations, *α*≈ 1 − (1 + *q*)^3^ ·*ν*_BSA_ · [BSA], with *q*_*s*_ = *δ/a* (*dashed*). Inset shows a magnified plot of the volume fraction in the range 0 mM *<* [BSA] *<* 2 mM. Curves are shown for PEG 4K (*yellow*), PEG 6K (*black*), PEG 8K (*blue*), PEG 20K (*red*), PEG 35K (*green*), and PEG 100K (*purple*).

The same crossover appears in the osmotic pressure (Fig. 4B). In the dilute regime, PEG obeys the van’t Hoff law for ideal solutions, Π = *RT*· [PEG]_*R*_. In the semidilute regime, the osmotic pressure is governed by the density of correlation blobs of volume 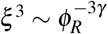. The interpolation between these regimes is

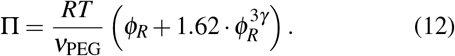

For *φ*_*R*_ ≫ 1, the second term dominates, giving

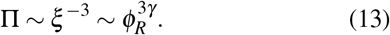

For PEG 20K, 35K, and 100K, the osmotic-pressure curves in the coexistence region follow the semidilute scaling approximation (Fig. 4B, *dashed*).

The free volume fraction along the binodal decreases with increasing BSA concentration and the accompanying increase in excluded volume (Fig. 4C). For sufficiently low BSA concentrations where overlap between depletion layers is negligible ([BSA]*<* 2 mM), the free volume decreases linearly with BSA concentration

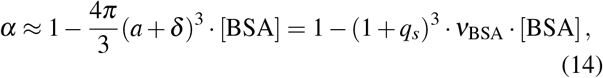

where *q*_*s*_ = *δ/a* and *ν*_BSA_ = 4*πa*^3^*N*_*A*_*/*3 is the molar volume of BSA.

We then evaluated the total colloidal osmotic pressure and BSA chemical potential along the best-fit GFVT binodals (Fig. 5). The total osmotic pressure is

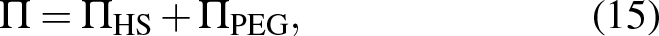

where Π_HS_ is the Carnahan-Starling hard-sphere contribution of BSA (Eq. 28) and Π_PEG_ is the depletant contribution (Eq. 30) [58]. The chemical potential of BSA can likewise be decomposed into hard-sphere and depletant contributions,

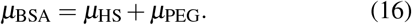

**FIG. 5.**
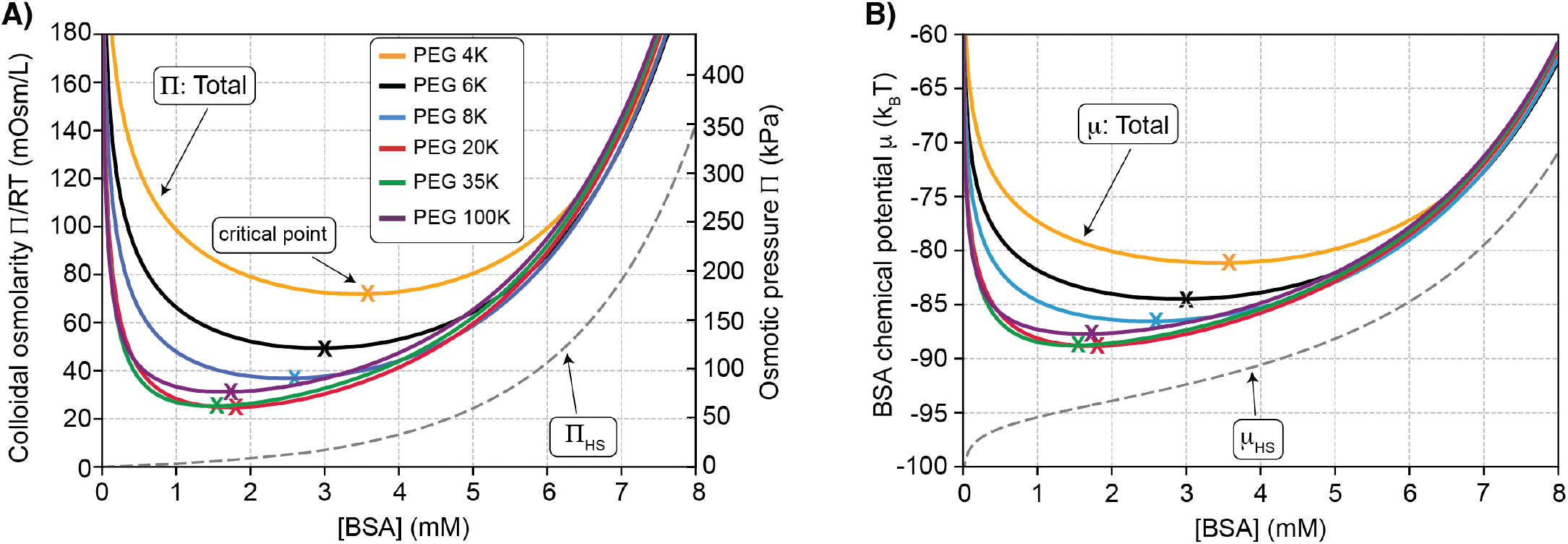
Osmotic pressure and BSA chemical potential along best-fit GFVT binodals. **A)** Total colloidal osmotic pressure, Π (Eq. 15, *solid*), osmotic pressure of the BSA hard-sphere fluid, Π_HS_ (Eq. 28, *dashed*) as a function of BSA concentration along the binodal. **B)** Total chemical potential of BSA in the presence of depletant, *µ* (Eq. 16, *solid*), and the contribution of the depletant-free BSA hard-sphere fluid *µ*_BSA_ (Eq. 29, *dashed*) as a function of BSA concentration along the binodal. GFVT predicted critical points (*crosses*). Curves are shown for PEG 4K (*yellow*), PEG 6K (*black*), PEG 8K (*blue*), PEG 20K (*red*), PEG 35K (*green*), and PEG 100K (*purple*).

At coexistence, the osmotic pressure and the BSA chemical potential are equal in the two phases (Eqs. 17 and 18), so the tie lines are horizontal in these thermodynamic representations. In the supernatant arm of the binodal, osmotic pressure and chemical potential are dominated by PEG, whereas in the dense condensate arm the BSA hard-sphere contribution becomes increasingly important. For PEG molecular weights of 20 kDa and above, both the osmotic pressure and the BSA chemical potential along the binodal become approximately independent of PEG molecular weight (Fig. 5). This molecular-weight invariance is the thermodynamic basis for the mass-concentration collapse observed in Fig. 3. The osmotic pressure due to PEG required for phase separation in this system is approximately 50-450 kPa (20-180 mOsm/L), exceeding reported colloidal osmotic pressures of 1-20 kPa in living cells [59, 60]. This comparison underscores that the BSA-PEG system is a controlled depletion model, not a direct measurement of cytoplasmic crowding, although cell extracts can be sufficiently crowded to induce BSA LLPS without added PEG [29].

### Polydispersity of the depletant PEG

Another practically important question is how polymer polydispersity affects the phase diagram. Biological crowders are often polydisperse (*e*.*g*. polysaccharides, heterogeneous protein mixtures), and even PEG may have a distribution of chain lengths. To test the simplest form of polydispersity, we prepared defined binary PEG mixtures and compared their BSA phase diagrams with single-component PEG phase diagrams (Fig. 6A). One mixture contained PEG 2K and PEG 8K with a weight-average molecular weight *M*_*w*_ = 7.3 kDa; the other contained PEG 2K and PEG 35K with *M*_*w*_ = 25 kDa. The measured binodal for the 2K + 8K mixture (*lilac symbols*) agreed with a GFVT binodal corresponding to monodisperse PEG 7.3K (*lilac curve*). Similarly, the 2K + 35K mixture (*light green symbols*) agreed well with the GFVT binodal for PEG 25K.

**FIG. 6.**
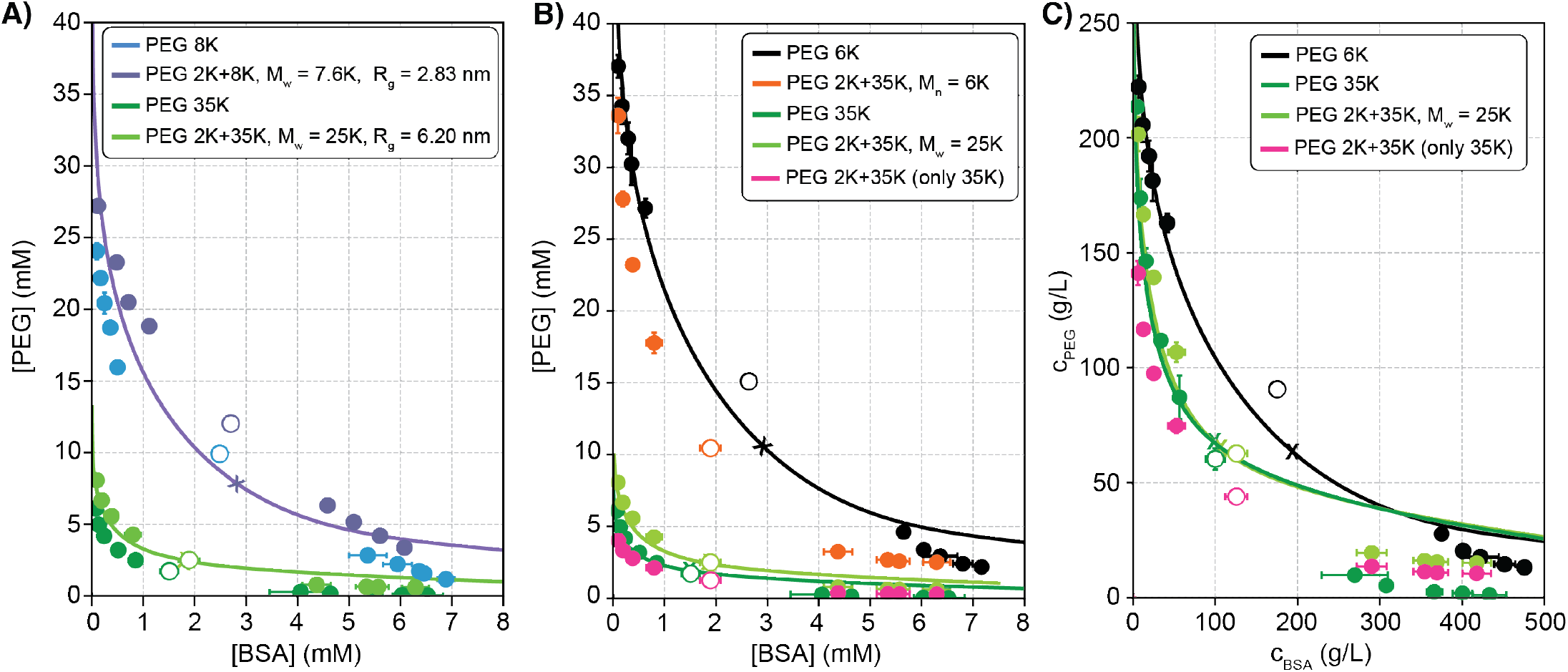
GFVT binodals correspond to the weight-average molecular weight of polydisperse PEG mixtures. **A)** Experimentally measured binodals (*circles*) and critical points (*open circles*), and GFVT binodals (*curves*) and critical points (*crosses*) for BSA in the presence of a mixture of PEG 2K and PEG 8K with a weight-average molecular weight of 7.3 kDa (q = 0.94, *lilac*), and a mixture of PEG 2K and PEG 35K with a weight-average molecular weight of 25 kDa (q = 2.07, *light green*). For comparison, experimental binodals and critical points in the presence of PEG 8K (*blue*) and PEG 35K (*green*) are shown. **B)** Experimentally measured binodals in the presence of the mixture of PEG 2K and PEG 35K plotted with respect to molar concentration assuming *M* = *M*_*n*_ = 6 kDa (*orange*), *M* = *M*_*w*_ = 25 kDa (*light green*). For comparison, binodals in the presence of monodisperse PEG 6 kDa (*black*) and PEG 35 kDa (*green*) are shown, along with the binodal of the 2K + 35 mixture assuming that only PEG 35 kDa contributes to crowding (*pink*). **C)** Experimentally measured binodals in the presence of the bimodally polydisperse mixture of PEG 2K and PEG 35K plotted with respect to mass concentration (*light green*). For comparison, binodals in the presence of approximately monodisperse (or unimodally polydisperse) PEG 6 kDa (*black*) and PEG 35 kDa (*green*) are shown, along with the binodal of the 2K + 35 mixture assuming that only PEG 35 kDa contributes to crowding (*pink*).

Because GFVT requires an effective polymer size, we tested whether the number-average molecular weight, *M*_*n*_, or the weight-average molecular weight, *M*_*w*_, provides the appropriate mapping. For the PEG 2K + 35K mixture, *M*_*w*_ = 25 kDa whereas *M*_*n*_ = 6 kDa. Plotting the experimental binodal in molar concentration units using *M*_*n*_ = 6 kDa (Fig. 6B, *orange*) did not agree with the GFVT binodal for PEG 6K (*black*). In contrast, plotting the data using *M*_*w*_ = 25 kDa agreed with the GFVT binodal for PEG 25K (Figs. 6A and B, *light green*).

Thus, for the defined binary mixtures tested here, PEG polydispersity is captured by an effective *R*_*g*_ corresponding to *M*_*w*_. The low-molecular-weight component nevertheless contributes to depletion. PEG 2K alone did not induce stable LLPS at room temperature, but in the PEG 2K + 35K mixture it shifted the binodal (Fig. 6B and C, *pink*) relative to a plot that counts only the PEG 35K concentration (*green*). This demonstrates that short chains can modulate the phase boundary even when they are individually too small to stabilize LLPS. In the semidilute regime, however, the scaling collapse implies that sensitivity to molecular-weight distribution weakens once the effective *M*_*w*_ is sufficiently high. Consistent with this expectation, the PEG 2K + 35K mixture with *M*_*w*_ = 25 kDa largely overlaps the pure PEG 35K binodal when plotted in mass concentration units (Fig. 6C).

## DISCUSSION

Measurements of both coexisting phases allow us to test how polymer scaling describes PEG-induced protein phase separation. In the dilute-chain regime, PEG molecular weight strongly affects the phase boundary because the depletion layer is set by the size of individual polymer coils. In the semidilute regime, the relevant depletion length is the polymer correlation length, and BSA-PEG phase diagrams approximately collapse in PEG mass concentration. GFVT connects these regimes by combining hard-sphere thermodynamics with scaling concepts of Flory-de Gennes polymer theory. The ovalbumin, MBP, and GB1 binodals extend this test for globular proteins beyond BSA (Fig. 2E). Their description uses independently specified protein radii and the PEG radius of gyration obtained from the BSA-PEG 35K binodal. Agreement without refitting either size parameter supports the transferability of the depletion-based model across the four globular proteins. Together with the binary-mixture measurements, these results test the model across changes in protein identity, protein size, and polymer composition.

The result clarifies the relation between conventional virial descriptions and polymer-scaling descriptions of PEG-induced protein phase separation. The effect of PEG on protein solubility has often been analyzed using virial expansions truncated at second order [61–65]. Such approaches can extract useful interaction parameters such as second virial coefficients from solubility data [30], but they become increasingly limited in crowded, non-ideal regimes where higher-order interactions are important. In the BSA concentration range studied here, the Carnahan-Starling equation of state for hard spheres accurately describes BSA up to 8 mM using a single parameter, the radius of the hard sphere *a*, but requires sixth-order virial coefficients to be approximated in this range [66]. Similarly, the equation of state of the depletant given by Eq. 12 deviates substantially from a second-order virial approximation over the concentration range relevant to phase separation. The semidilute collapse therefore reflects polymer collective behavior rather than a simple pairwise correction to protein solubility.

The full binodal measurements are also important because the supernatant arm alone does not determine the complete thermodynamics of the two-phase system. Protein solubility curves are often interpreted by assuming that the protein chemical potential in the dense phase is effectively constant, or that PEG does not appreciably partition into the condensate [38, 62, 63]. The measured condensate arms show finite PEG partitioning that approaches unity near the critical point (Fig. 2A), consistent with prior observations that polymer partitioning can remain important even near solid precipitation boundaries [67]. Direct tie-line measurements therefore provide constraints that are not available from cloud-point or solubility measurements alone.

The semidilute collapse concerns changes in PEG molecular weight at fixed protein identity. It does not imply that different proteins share a common binodal in mass-concentration coordinates. The collapse is expected when the crowder is a neutral or weakly interacting polymeric depletant, the relevant coexistence region lies above the polymer overlap concentration, and protein-polymer attractions are negligible. These conditions should apply most directly to disordered polymeric crowders that are described by Flory-de Gennes scaling. Other architectures, including branched polymers such as dextran and Ficoll, may show analogous behavior but with architecture-dependent scaling exponents [68]. Conversely, attractive protein-polymer interactions can shift or disrupt the depletion-only collapse. Such interactions may contribute to observations that dextran does not induce BSA LLPS under conditions where PEG does [69], and that PEG-lysozyme cloud points depend on PEG size even though the depletion-only GFVT model contains no explicit temperature dependence [70, 71].

Polydispersity introduces a second practical boundary condition. For the binary PEG mixtures tested here, the phase diagram was approximated by a monodisperse PEG with the same weight-average molecular weight (Fig. 6A and B). This observation is useful because commercial and biological polymers are rarely monodisperse. The measured binodal for bi-modally polydisperse PEG 2K + 35K (Fig. 6C, *light green*) is slightly narrower than the PEG 35K binodal (*green*). This agrees with previous observations of the PEG-dextran system, where polydispersity was found to narrow the two-phase region without affecting the critical point due to partitioning of shorter dextran chains to the PEG-rich phase and the two-phase interface [72]. However, GFVT overestimates the PEG concentration in the condensate phase when plotting the phase diagram with respect to mass concentration (Fig. 3A).

Because polydispersity tends to narrow two-phase regions whereas the measured binodals are wider than the GFVT prediction, this discrepancy is unlikely to be explained by the finite polydispersity of our PEG stocks [72]. It may instead reflect protein-specific physics omitted from the hard-sphere representation, such as BSA conformational changes or weak protein–polymer interactions. Together, these results establish semidilute polymer scaling as a practical metric for comparing PEG-induced crowding, while clarifying where additional protein-specific physics must be included.

## CONCLUSION

We measured full BSA-PEG liquid-liquid phase diagrams and showed that GFVT quantitatively describes the binodal and tie lines using the PEG radius of gyration as the polymer-size parameter. Deviations near critical points are consistent with the expected limitations of mean-field theory. In PEG 35K, GFVT also describes the binodals of ovalbumin, MBP, and GB1 using independently specified protein radii and the PEG radius of gyration calibrated against BSA. Together with BSA, these measurements test the model across proteins spanning 7.2–66.4 kDa. The BSA measurements identify a crossover from phase boundaries that depend strongly on PEG chain size to an approximately common phase diagram in mass concentration units for PEG molecular weights of 20 kDa and above. For the binary mixtures tested, the binodals are approximated using an effective polymer size corresponding to the weight-average molecular weight. These results identify conditions under which PEG mass concentration permits comparisons of depletion-induced phase separation and test the applicability of GFVT across several globular proteins. Future work should extend this framework to include hydration, electrostatics, polymer architecture, and weak protein-crowder interactions. These additions would help separate depletion from specific solvation effects in more complex condensate-forming systems such as cytoplasm or synthetic cells [10, 69, 73].

## ACKNOWLEDGEMENTS

This research was supported by an AD416 faculty research grant to AAR. Experiments were carried out using the Core Technology Platforms resources at New York University Abu Dhabi. We thank Remco Tuinier and Robert W. Style for helpful suggestions and insightful discussions. ChatGPT 5.6 was used to aid writing technically complicated sentences with the prompt “revise to enhance clarity”.

## AUTHOR CONTRIBUTIONS

A.A.R. conceived and supervised the project. A.A.R., M.A., M.T.W., and H.O.S. performed the experiments. A.A.R. and A.I. implemented the GFVT model and interpreted the experiments. A.A.R. wrote the paper with input from all authors.

## MATERIALS AND METHODS

### Materials

Polyethylene glycol 4000 Da (PEG 4K) (A16161.0B), PEG 6K (A17541.30), PEG 8K (043443.22), PEG 20K (81300), PEG 35K (94646), PEG 100K (181986), bovine serum albumin (BSA) (A7638), ovalbumin (A5503) potassium phosphate dibasic (P8281), potassium phosphate monobasic (P5379), potassium chloride (KCl) (529552), sodium chloride (746398), Tris-HCl (T5941), Tris-base (T1503), deuterium oxide (D_2_O) (151882), N,N-dimethylformamide (DMF) (103053), BL21 Star (DE3) pLysS cells (C602003), and Ni-NTA agarose resin (88222) were purchased from either Sigma-Aldrich or Fisher Scientific. All water used was Milli-Q.

### Stock Preparation

Stock solutions of polyethylene glycol were prepared by dissolving the PEG variant in Milli-Q water. The PEG stock concentrations were confirmed using proton nuclear magnetic resonance (^1^H NMR) spectroscopy [74]. NMR samples were prepared by mixing a fixed volume of the PEG solution (100 µL) with dimethylformamide (DMF; 100 µL, 13.7 M), which served as a calibration standard, and deuterium oxide (D_2_O; 400 µL). The mixture was vortexed and transferred into a disposable 5 mm NMR tube (Wilmad WG10007). ^1^H NMR spectra were acquired on a 500 MHz spectrometer (Bruker Avance 500 MHz) at 298 K and averaged 16 times. The data were analyzed using MestReNova software (Mestrelab Research). The peaks at 3.02 ppm and 2.86 ppm correspond to the two methyl groups in DMF. Accordingly, the sum of the integrals of the two peaks corresponds to the signal from six protons per DMF molecule and serves as a calibration standard given the known DMF concentration in the solution. The PEG peak occurs at 3.71 ppm and its integral corresponds to the contribution from~ MW*/*11 protons per PEG molecule, where MW is the molecular weight of PEG in Da. The PEG concentrations were calculated based on the relative integral of the 3.71 ppm peak to the sum of the integrals of the two DMF peaks at 3.02 ppm and 2.86 ppm.

The BSA stock solution with a target concentration of 7.5 mM (500 mg/ml) was prepared by mixing the appropriate amount of powdered BSA and Milli-Q water. The actual concentration was confirmed by measuring the absorption intensity at the 280 nm peak with a UV-Vis spectrophotometer (NanoDrop One/OneC UV-Vis Spectrophotometer) using a quartz cuvette (UQ-124, Portmann Technologies), and using the experimentally determined BSA extinction coefficient of *ε* = 41, 190 M^−1^cm^−1^. The ovalbumin stock solution was prepared by dialyzing 5 g of ovalbumin against 40 mM Tris (pH 7.5) and 100 mM NaCl, followed by concentration to approximately 7.5 mM (332 mg/mL) using Amicon filter with 10 KDa molecular weight cutoff (UFC901024).

A 4X potassium phosphate buffer (KP) stock solution with a final KCl concentration of 800 mM and a final potassium phosphate concentration of 400 mM was prepared by mixing solutions of 800 mM KCl and 400 mM monobasic potassium phosphate and 400 mM dibasic potassium phosphate at a ratio that yielded pH 7. The pH of the solution was measured using a pH meter (Thermo Scientific Orion Star A211). A 4X Tris buffer (TB) stock solution, containing 200 mM Tris and 400 mM NaCl for GB1 and ovalbumin, or 800 mM NaCl for MBP, was prepared by diluting a pre-made 1 M Tris stock (pH 7.0), prepared from Tris base and Tris-HCl, and adding NaCl to the indicated final concentration. The pH of the final solution was verified and adjusted to 7.0 using NaOH, when necessary.

### Protein Expression and Purification

Maltose-binding protein (MBP) construct was derived from Addgene plasmid #135464 [75] by introducing a stop codon immediately downstream of the MBP coding sequence. The resulting protein has a molecular weight of 41.7 kDa. The GB1 construct was purchased from Addgene (plasmid #133438)[76]. MBP and GB1 were expressed in *Escherichia coli* BL21 Star (DE3) pLysS cells. 10 L cultures were grown in Luria–Bertani (LB) medium at 37 °C until reaching an optical density at 600 nm of approximately 0.6. Protein expression was induced with 0.5 mM isopropyl *β*-D-1-thiogalactopyranoside (IPTG), and the cultures were incubated for an additional 4 h at 37 °C for MBP or 20 h at 20 °C for GB1. Cell pellets were resuspended in lysis buffer containing 50 mM Tris-HCl (pH 7.5), 500 mM NaCl, 10 mM imidazole, 1 mg/mL lysozyme, 0.25% Triton X-100, 0.25 mM phenylmethylsulfonyl fluoride (PMSF), and one tablet of cOmplete, EDTA-free Protease Inhibitor Cocktail, followed by sonication for 3 min. The lysates were clarified by centrifugation at 50,000 ×g for 1 h at 4 °C. MBP and GB1 were purified using Ni-NTA agarose resin. Imidazole-eluted fractions containing MBP or GB1 were pooled and buffer-exchanged into imidazole-free buffer containing 50 mM Tris-HCl (pH 7.5) and 400 mM NaCl for MBP, or 50 mM Tris-HCl (pH 7.5) and 100 mM NaCl for GB1. The pooled MBP and GB1 fractions were then concentrated to final protein concentrations of 9 mM (387 mg/mL) and 54 mM (388 mg/mL) using Amicon filters with molecular weight cutoffs of 10 KDa and 3 KDa (UFC900308), respectively.

### Condensate Preparation

To form condensates, BSA, Milli-Q water, 4X KP buffer, and PEG were added to an Eppendorf tube in that order. For ovalbumin, MBP, and GB1 condensates, 4X TB was used instead of 4X KP buffer. The samples were kept at room temperature (22 °C) throughout the sample preparation and subsequent measurements. The mixture was gently vortexed to ensure thorough mixing, resulting in a turbid solution. The dense protein-rich condensate phase was then separated from the PEG-rich supernatant phase by centrifugation at 22 °C for 5 minutes at 3,000 × g.

### Measurement of Binodal

To determine the two points defining a tie line, the protein and PEG concentrations of each phase were measured for every condensate suspension prepared. BSA concentrations were determined by measuring the UV-Vis absorbance of each sample at 280 nm using a quartz cuvette as described above. For ovalnumin, MBP, and GB1, absorbance at 280 nm was measured using the pedestal of the Nanodrop (NanoDrop One/OneC UV-Vis Spectrophotometer). PEG concentrations were measured using ^1^H NMR, as described above in the stock preparations. For full binodal measurements, this procedure was repeated at progressively lower concentrations of protein and PEG in the final mixture until the critical point was reached.

### Measurement of Critical Point

A condensate suspension was prepared with final BSA and PEG concentrations near the anticipated critical point. After centrifugation, equal volumes of each phase were collected and transferred to a new tube, then diluted with a small volume of KP buffer to precisely approach the critical point. This process of centrifugation, phase transfer, and incremental dilution was repeated until the solution remained turbid, but the compositions and densities of the two phases became too similar to allow further separation by centrifugation. The critical point was then determined by measuring the BSA and PEG concentrations as described above.

### Phase Diagram Calculation Using GFVT

We used generalized free volume theory (GFVT) to calculate binodals, critical points, and spinodals, representing proteins as colloidal hard spheres of radius *a* and PEG as a semi-flexible polymer with radius of gyration *R*_*g*_ [18, 57]. For the BSA-PEG series, the protein radius was fixed at *a* = 3.0 nm and *R*_*g*_ was fitted to the BSA binodal for each PEG molecular weight. For the additional protein binodals in PEG 35K, the protein radii were specified independently of the binodal fits: *a* = 2.5 nm for MBP, *a* = 2.7 nm for ovalbumin, and *a* = 1.5 nm for GB1. These calculations retained the PEG radius of gyration calibrated against BSA-PEG 35K, without fitting either size parameter to the additional protein binodals.

GFVT yields expressions for the colloidal osmotic pressure, Π, and the chemical potential of the hard spheres, *µ*_BSA_, as a function of hard sphere volume fraction, *η* = 4*πa*^3^*n*_*η*_ */*3, and the relative concentration of depletant, 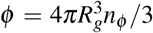, where *n*_*η*_ and *n*_*φ*_ are the number densities of colloid and depletant, respectively. It is convenient to use the dimensionless pressure 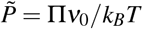 and dimensionless chemical potential 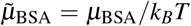, where *ν*_0_ is the volume of the colloidal hard sphere, *k*_*B*_ is the Boltzmann constant, and *T* is the absolute temperature. At the binodal, the following equations must be satisfied:

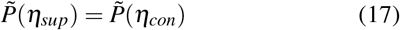

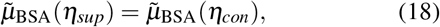

where *η*_*sup*_ and *η*_*con*_ are the colloid (BSA) volume fractions in the supernatant and condensate phases, respectively. In practice, the system is assumed to be in equilibrium with a reservoir of depletant with a depletant concentration *φ*_*R*_. Both phases are in equilibrium with the same reservoir and therefore Eqs. 17 and 18 are solved by iterating *φ*_*R*_ over a fixed range and then back-calculating *φ* from *φ*_*R*_ for each phase using the respective free volume fraction, *α*(*η*):

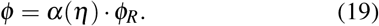

The free volume fraction obtained from scaled particle theory is a function of both the BSA volume fraction, *η*, and the PEG concentration in the reservoir, *φ*_*R*_:

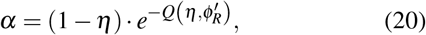

where

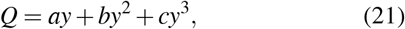

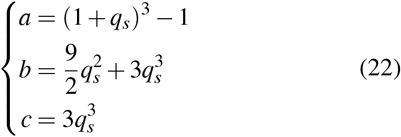

and

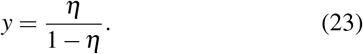

It is helpful to abbreviate *∂*^*n*^*X/∂y*^*n*^ as *X*_*n*_ and to define

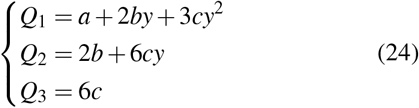

and

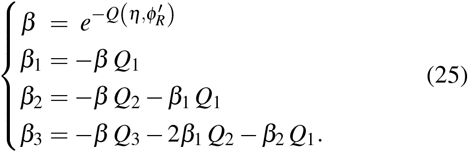

Both 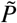 and 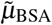 can be written as a sum of contributions from the hard-spheres and the depletant:

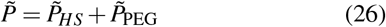

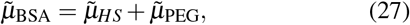

where 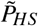 and 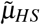 are the dimensionless osmotic pressure and chemical potential of the hard sphere fluid in the absence of depletant, and 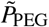 and 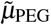 are the corrections to the dimensionless osmotic pressure and chemical potential, respectively, due to the addition of depletant. We use the Carnahan-Starling expression for the dimensionless pressure of a hard sphere fluid [58],

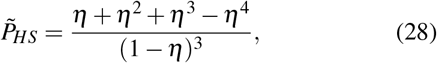

and the corresponding expression for the dimensionless chemical potential

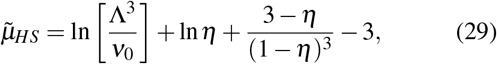

where 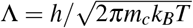 is the de Broglie wavelength for a colloidal particle with mass *m*_*c*_. Note that the first term involving the de Broglie wavelength cancels out when solving Eq. 18. The contributions of the depletant are given by

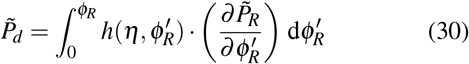

and

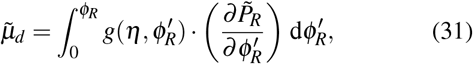

where 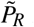 is the dimensionless osmotic pressure of the depletant in the reservoir. The functions *g* and *h* are defined as follows:

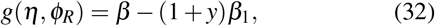

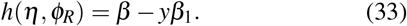

Here, we use a GFVT formulation in which the PEG depletant is treated as a semi-flexible polymer in good solvent. The depletion thickness, *q*_*s*_, then becomes

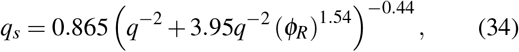

with *q* = *R*_*g*_*/a*, where *a* is the hard-sphere radius of the protein and *R*_*g*_ is the radius of gyration of PEG. In good solvent, the osmotic compressibility is given by

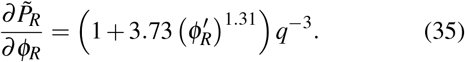

Solving Eqs. 17 and 18 for a given *φ*_*R*_ yields the volume fractions of protein in the supernatant and condensate phases, *η*_sup_ and *η*_con_, respectively. The relative depletant concentration in each phase is related to the relative concentration in the reservoir via

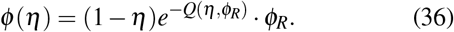

The molar concentrations of BSA and PEG in each phase, in mmol/L, are then given by

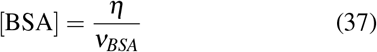

and

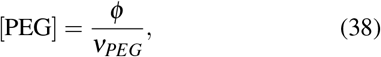

where *ν*_*BSA*_ and *ν*_*PEG*_ are the molar volumes of BSA and PEG, respectively. We use values that are self-consistent with the model:

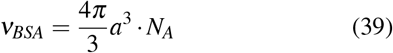

and

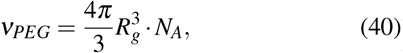

where *N*_*A*_ is Avogadro’s constant.

The critical point, (*η, φ*_*R*_)_cp_, is calculated by simultaneously solving

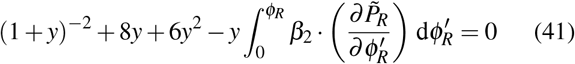

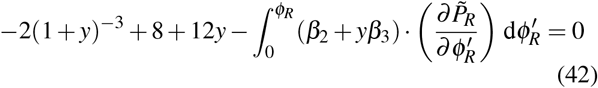

for *y*. The depletant concentration in the reservoir at the critical point, (*φ*_*R*_)_cp_, is a suitable lower limit for the range of values for which the binodal calculation (Eqs. 17 and 18) should be performed, *e*.*g*., *φ*_*R*_ ∈ (*φ*_*R*_)_cp_, 1.

### Analytical Approximation to Binodal

To compute the analytical approximation to the binodal, we first calculate the triple point, (*η*_*sup*_, *η*_*cond*_, *η*_*sol*_, *φ*_*R*_)_tp_, by equating the chemical potentials and osmotic pressures in the supernatant, condensate, and putative solid phases:

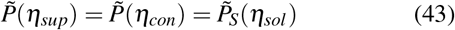

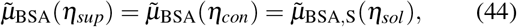

where

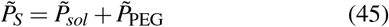

with

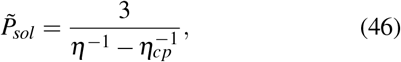

and

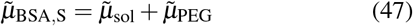

with

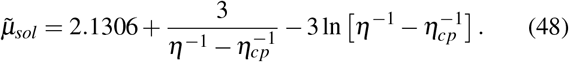

The analytical binodal was calculated for the supernatant and condensate arms separately by interpolating between the critical point (*η*_*c*_, *φ*_*R*_)_crit_ and the triple points, (*η*_sup_, *φ*_*R*_) _tp_, and (*η*_con_, *φ*_*R*_)_tp_, as described in Fleer et al. [57]. For both arms, we performed the computation by iterating *φ*_*R*_ ∈ [(*φ*_*R*_)_*cp*_, (*φ*_*R*_)_*t p*_]. For the supernatant arm of the binodal, we define the relative depletion thickness as a function of *φ*_*R*_:

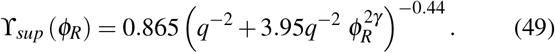

For the condensate arm of the binodal, we define the osmotic pressure of PEG in the reservoir as a function of *φ*_*R*_:

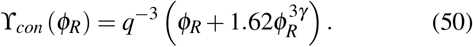

The supernatant arm of the binodal for *φ*_*R*_ ∈ (*φ*_*R*_)_*cp*_, (*φ*_*R*_)_*t p*_ then becomes

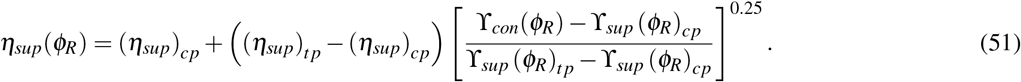

Similarly, the condensate arm of the binodal is constructed using

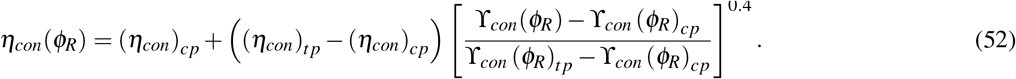

Finally, the relative PEG concentrations in the reservoir (*i*.*e*. in terms of the free volume) are converted to relative PEG concentrations in the system (*i*.*e*. in terms of the total volume) (Eq. 19):

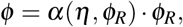

where *α* is calculated using Eqs. 20-23 and Eq. 34.

### Statistics and Reproducibility

Phase diagrams shown in Fig. 1B, Fig. 2A, and Fig. 6 (with the exception of Fig. 2E and PEG 2K+8K in Fig. 6A) were measured in triplicate, with all data points possessing both horizontal and vertical error bars denoting standard deviations. Some error bars are not visible due to small size. The measurements for ovalbumin, MBP, and GB1 shown in Fig. 2E and the measurements for PEG 2K+8K shown in Fig. 6A were repeated once.

## Data Availability

The data that support the findings of this article are available at https://doi.org/10.5281/zenodo.19865984.

## Code Availability

The codes used in this study are available at https://doi.org/10.5281/zenodo.19865984.

## References

[1] Y. Shin and C. P. Brangwynne, Liquid phase condensation in cell physiology and disease, Science 357, eaaf4382 (2017), https://www.science.org/doi/pdf/10.1126/science.aaf4382.

[2] S. F. Banani, H. O. Lee, A. A. Hyman, and M. K. Rosen, Biomolecular condensates: organizers of cellular biochemistry, Nature Reviews Molecular Cell Biology 18, 285 (2017).

[3] F. C. Keber, T. Nguyen, A. Mariossi, C. P. Brangwynne, and M. Wühr, Evidence for widespread cytoplasmic structuring into mesoscale condensates., Nature cell biology (2024).

[4] C. P. Brangwynne, C. R. Eckmann, D. S. Courson, A. Rybarska, C. Hoege, J. Gharakhani, F. Jülicher, and A. A. Hyman, Germline p granules are liquid droplets that localize by controlled dissolution/condensation, Science 324, 1729 (2009), https://www.science.org/doi/pdf/10.1126/science.1172046.

[5] M. Feric, N. Vaidya, T. S. Harmon, D. M. Mitrea, L. Zhu, T. M. Richardson, R. W. Kriwacki, R. V. Pappu, and C. P. Brangwynne, Coexisting liquid phases underlie nucleolar subcompartments, Cell 165, 1686 (2016).

[6] A. Molliex, J. Temirov, J. Lee, M. Coughlin, A. P. Kanagaraj, H. J. Kim, T. Mittag, and J. P. Taylor, Phase separation by low complexity domains promotes stress granule assembly and drives pathological fibrillization, Cell 163, 123 (2015).

[7] M. Zeng, Y. Shang, Y. Araki, T. Guo, R. L. Huganir, and M. Zhang, Phase transition in postsynaptic densities underlies formation of synaptic complexes and synaptic plasticity, Cell 166, 1163 (2016).

[8] T. Kaur, I. Alshareedah, W. Wang, J. A. Ngo, M. M. Moosa, and P. R. Banerjee, Molecular crowding tunes material states of ribonucleoprotein condensates, Biomolecules 9 (2018).

[9] M. D. Crabtree, J. Holland, A. S. Pillai, P. Kompella, L. Babl, N. N. Turner, J. T. Eaton, G. K. A. Hochberg, D. G. A. L. Aarts, C. Redfield, A. J. Baldwin, and T. J. Nott, Ion binding with charge inversion combined with screening modulates dead box helicase phase transitions, Cell Reports 42 (2023).

[10] X. Xu, A. A. Rebane, L. R. Julià, K. A. Rosowski, E. R. Dufresne, and F. Stellacci, Amino acids modulate liquid–liquid phase separation in vitro and in vivo by regulating protein–protein interactions, Proceedings of the National Academy of Sciences of the United States of America 121 (2024).

[11] X. Xu, M. H. I. van Haren, I. B. A. Smokers, B. S. Visser, P. B. White, R. S. Jansen, and E. Spruijt, Amino acids bind to phase-separating proteins and modulate biomolecular condensate stability and dynamics, Nature Communications 16 (2025).

[12] E. Jambon-Puillet, A. Testa, C. Lorenz, R. W. Style, A. A. Rebane, and E. R. Dufresne, Phase-separated droplets swim to their dissolution, Nature Communications 15 (2023).

[13] S. B. Zimmerman and S. O. Trach, Estimation of macro-molecule concentrations and excluded volume effects for the cytoplasm of escherichia coli, Journal of Molecular Biology 222, 599 (1991).

[14] R. Ellis, Macromolecular crowding: obvious but underappreciated., Trends in biochemical sciences 26 10, 597 (2001).

[15] H. Zhou, G. Rivas, and A. P. Minton, Macromolecular crowding and confinement: biochemical, biophysical, and potential physiological consequences., Annual review of biophysics 37, 375 (2008).

[16] R. Hancock, Crowding, entropic forces, and confinement: Crucial factors for structures and functions in the cell nucleus, Biochemistry (Moscow) 83, 326 (2018).

[17] C. Alfano, Y. Fichou, K. Huber, M. Weiss, E. Spruijt, S. Ebbinghaus, G. De Luca, M. A. Morando, V. Vetri, P. A. Temussi, and A. Pastore, Molecular crowding: The history and development of a scientific paradigm, Chemical Reviews 124, 3186 (2024), pMID: 38466779, 10.1021/acs.chemrev.3c00615.

[18] H. N. W. Lekkerkerker and R. Tuinier, Colloids and the depletion interaction, Lecture Notes in Physics (2011).

[19] S. Asakura and F. Oosawa, On interaction between two bodies immersed in a solution of macromolecules, Journal of Chemical Physics 22, 1255 (1954).

[20] A. Vrij, Polymers at interfaces and the interactions in colloidal dispersions, Pure and Applied Chemistry 48, 471 (1976).

[21] F. Zosel, A. Soranno, K. J. Buholzer, D. Nettels, and B. Schuler, Depletion interactions modulate the binding between disordered proteins in crowded environments, Proceedings of the National Academy of Sciences 117, 13480 (2020), https://www.pnas.org/doi/pdf/10.1073/pnas.1921617117.

[22] Y. C. Kim and J. Mittal, Crowding induced entropy-enthalpy compensation in protein association equilibria, Phys. Rev. Lett. 110, 208102 (2013).

[23] G. Wolfisberg, J. Agudo-Canalejo, P. C. Bittmann, E. R. Dufresne, R. W. Style, and A. A. Rebane, Dynamic shape modulation of deflated and adhered lipid vesicles, Journal of the American Chemical Society 147, 38720 (2025), pMID: 41071582, 10.1021/jacs.5c13925.

[24] A. M. Kulkarni, A. P. Chatterjee, K. S. Schweizer, and C. F. Zukoski, Effects of polyethylene glycol on protein interactions, Journal of Chemical Physics 113, 9863 (2000).

[25] O. Annunziata, N. Asherie, A. Lomakin, J. Pande, O. Ogun, and G. B. Benedek, Effect of polyethylene glycol on the liquid–liquid phase transition in aqueous protein solutions, Proceedings of the National Academy of Sciences of the United States of America 99, 14165 (2002).

[26] O. Annunziata, O. Ogun, and G. B. Benedek, Observation of liquid–liquid phase separation for eye lens γs-crystallin, Proceedings of the National Academy of Sciences of the United States of America 100, 970 (2003).

[27] R. Tuinier and Á. González García, Associative and segregative liquid-liquid phase separation in macromolecular solutions, Newton 2, 100550 (2026).

[28] S. Alberti, S. Saha, J. B. Woodruff, T. M. Franzmann, J. Wang, and A. A. Hyman, A user’s guide for phase separation assays with purified proteins, Journal of Molecular Biology 430, 4806 (2018).

[29] M. Poudyal, K. Patel, L. Gadhe, A. S. Sawner, P. Kadu, D. Datta, S. Mukherjee, S. Ray, A. Navalkar, S. Maiti, D. Chatterjee, J. Devi, R. Bera, N. Gahlot, J. Joseph, R. Padinhateeri, and S. K. Maji, Intermolecular interactions underlie protein/peptide phase separation irrespective of sequence and structure at crowded milieu, Nature Communications 14 (2023).

[30] G. Chauhan, A. Bremer, F. Dar, T. Mittag, and R. V. Pappu, Crowder titrations enable the quantification of driving forces for macromolecular phase separation., Biophysical journal (2023).

[31] A. A. Bhattacharya, T. Grune, and S. Curry, Crystallographic analysis reveals common modes of binding of medium and long-chain fatty acids to human serum albumin., Journal of molecular biology 303 5, 721 (2000).

[32] A. Testa, M. Dindo, A. A. Rebane, B. Nasouri, R. W. Style, R. Golestanian, E. R. Dufresne, and P. Laurino, Sustained enzymatic activity and flow in crowded protein droplets, Nature Communications 12 (2021).

[33] R. Koningsveld and A. J. Staverman, Liquid–liquid phase separation in multicomponent polymer solutions. ii. the critical state, Journal of Polymer Science Part A-2: Polymer Physics 6, 325 (1968).

[34] Y. Liu, R. Lipowsky, and R. Dimova, Concentration dependence of the interfacial tension for aqueous two-phase polymer solutions of dextran and polyethylene glycol., Langmuir : the ACS journal of surfaces and colloids 28 8, 3831 (2012).

[35] R. Tuinier, P. A. Smith, W. C. K. Poon, S. U. Egelhaaf, D. G. A. L. Aarts, H. N. W. Lekkerkerker, and G. J. Fleer, engPhase diagram for a mixture of colloids and polymers with equal size, Europhysics letters 82, 68002 (2008).

[36] I. C. Sanchez, Critical amplitude scaling laws for polymer solutions, The Journal of Physical Chemistry 93, 6983 (1989).

[37] W. C. K. Poon, The physics of a model colloid–polymer mixture, Journal of Physics: Condensed Matter 14, R859 (2002).

[38] Y. Wang and O. Annunziata, Comparison between protein-polyethylene glycol (peg) interactions and the effect of peg on proteinprotein interactions using the liquidliquid phase transition, The Journal of Physical Chemistry B 111, 1222 (2007), pMID: 17266278, 10.1021/jp065608u.

[39] K. Devanand and J. C. Selser, Asymptotic behavior and long-range interactions in aqueous solutions of poly(ethylene oxide), Macromolecules 24, 5943 (1991).

[40] L. Almásy, O. P. Artykulnyi, V. I. Petrenko, O. I. Ivankov, L. A. Bulavin, M. Yan, and V. M. Haramus, Structure and intermolecular interactions in aqueous solutions of polyethylene glycol, Molecules 27 (2022).

[41] A. R. Denton, Effective interactions and volume energies in charge-stabilized colloidal suspensions, Journal of Physics: Condensed Matter 11, 10061 (1999).

[42] C. Gögelein and R. Tuinier, Phase behaviour of a dispersion of charge-stabilised colloidal spheres with added non-adsorbing interacting polymer chains, The European Physical Journal E 27, 171 (2008).

[43] J. Song, R. P. Tas, M. C. M. Martens, M. V. Ritten, H. Wu, E. R. Jones, J. G. J. L. Lebouille, M. Vis, I. K. Voets, and R. Tuinier, Freezing-mediated formation of supraproteins using depletion forces., Journal of colloid and interface science 665, 622 (2024).

[44] M. C. Martens, Z. Han, R. V. Freire, S. Huisman, I. K. Voets, T. P. Knowles, R. Tuinier, and N. A. Erkamp, Protein self-assembly in crowded environments, Journal of Colloid and Interface Science 727, 141573 (2027).

[45] T. Matsumoto and H. Inoue, Association state, overall structure, and surface roughness of native ovalbumin molecules in aqueous solutions at various ionic concentrations, Journal of Colloid and Interface Science 160, 105 (1993).

[46] P. G. Telmer and B. H. Shilton, Insights into the conformational equilibria of maltose-binding protein by analysis of high affinity mutants*, Journal of Biological Chemistry 278, 34555 (2003).

[47] S. Pradhan, R. Rath, and M. Biswas, Gb1 dimerization in crowders: A multiple resolution approach, Journal of Chemical Information and Modeling 63, 1570 (2023), https://pubs.acs.org/jcisd8/article-pdf/63/5/1570/9515836/ci3c00012.pdf.

[48] F. C. Cheong, S. Y. Lee, S. Bais, S. Khanal, and S. Saurabh, Holographic fingerprinting reveals oligomer-driven phase separation in bovine serum albumin, Chemical & Biomedical Imaging 0, null (0), 10.1021/cbmi.5c00258.

[49] P. G. de Gennes, Scaling concepts in polymer physics (1979).

[50] H. Hofmann, A. Soranno, A. Borgia, K. Gast, D. Nettels, and B. Schuler, Polymer scaling laws of unfolded and intrinsically disordered proteins quantified with single-molecule spectroscopy, Proceedings of the National Academy of Sciences 109, 16155 (2012), https://www.pnas.org/doi/pdf/10.1073/pnas.1207719109.

[51] G. J. Fleer, A. M. Skvortsov, and R. Tuinier, A simple relation for the concentration dependence of osmotic pressure and depletion thickness in polymer solutions, Macromolecular Theory and Simulations 16, 531 (2007).

[52] G. J. Fleer and R. Tuinier, Analytical phase diagram for colloid-polymer mixtures, Phys. Rev. E 76, 041802 (2007).

[53] P. G. Bolhuis, E. J. Meijer, and A. A. Louis, Colloid-polymer mixtures in the protein limit, Phys. Rev. Lett. 90, 068304 (2003).

[54] K. J. Mutch, J. S. van Duijneveldt, J. Eastoe, I. Grillo, and R. K. Heenan, Testing the scaling behavior of microemulsionpolymer mixtures, Langmuir 25, 3944 (2009), pMID: 19007255, 10.1021/la802488f.

[55] K. J. Mutch, J. S. van Duijneveldt, J. Eastoe, I. Grillo, and R. K. Heenan, Scaling the structure factors of protein limit colloidpolymer mixtures, Langmuir 26, 1630 (2010), pMID: 19761255, 10.1021/la902508z.

[56] G. J. Fleer, A. M. Skvortsov, and R. Tuinier, Mean-field equation for the depletion thickness, Macromolecules 36, 7857 (2003), 10.1021/ma0345145.

[57] G. J. Fleer and R. Tuinier, Analytical phase diagrams for colloids and non-adsorbing polymer., Advances in colloid and interface science 143 1-2, 1 (2008).

[58] N. F. Carnahan and K. E. Starling, Equation of state for nonattracting rigid spheres, The Journal of Chemical Physics 51, 635 (1969), https://pubs.aip.org/aip/jcp/article-pdf/51/2/635/18863564/6351online.pdf.

[59] T. J. Mitchison and T. J. Mitchison, Colloid osmotic parameterization and measurement of subcellular crowding, Molecular Biology of the Cell 30, 173 (2019).

[60] B. Liu, C. Åberg, F. J. van Eerden, S. J. Marrink, B. Poolman, and A. J. Boersma, Design and properties of genetically encoded probes for sensing macromolecular crowding., Biophysical journal 112 9, 1929 (2017).

[61] E. Edmond and A. G. Ogston, An approach to the study of phase separation in ternary aqueous systems., The Biochemical journal 109 4, 569 (1968).

[62] D. H. Atha and K. C. Ingham, Mechanism of precipitation of proteins by polyethylene glycols. analysis in terms of excluded volume., The Journal of biological chemistry 256 23, 12108 (1981).

[63] S.-L. Sim, T. He, A. Tscheliessnig, M. Mueller, R. B. H. Tan, and A. Jungbauer, Protein precipitation by polyethylene glycol: a generalized model based on hydrodynamic radius., Journal of biotechnology 157 2, 315 (2012).

[64] A. P. Minton, Simple calculation of phase diagrams for liquid–liquid phase separation in solutions of two macromolecular solute species, The Journal of Physical Chemistry B 124, 2363 (2020), pMID: 32118433, 10.1021/acs.jpcb.0c00402.

[65] A. Bot and P. Venema, Phase behavior of ternary polymer mixtures in a common solvent, ACS Omega 8, 28387 (2023), 10.1021/acsomega.3c02604.

[66] A. P. Minton, The effective hard particle model provides a simple, robust, and broadly applicable description of nonideal behavior in concentrated solutions of bovine serum albumin and other nonassociating proteins, Journal of Pharmaceutical Sciences 96, 3466 (2007).

[67] H. N. W. Lekkerkerker, W. C.-K. Poon, P. N. Pusey, A. Stroobants, and P. B. Warren, Phase behaviour of colloid + polymer mixtures, Europhysics Letters 20, 559 (1992).

[68] J. A. M. Smit, J. A. P. P. Van Dijk, M. G. Mennen, and M. Daoud, Polymer size exponents of branched dextrans, Macromolecules 25, 3585 (1992), 10.1021/ma00039a044.

[69] J. L. Watson, E. Seinkmane, C. T. Styles, A. Mihut, L. K. Krüger, K. E. McNally, V. J. Planelles-Herrero, M. Dudek, P. M. McCall, S. Barbiero, M. V. Oever, S. Y. Peak-Chew, B. T. Porebski, A. Zeng, N. M. Rzechorzek, D. C. S. Wong, A. D. Beale, A. Stangherlin, M. Riggi, J. H. Iwasa, J. Morf, C. Miliotis, A. Guna, A. J. Inglis, J. Brugués, R. M. Voorhees, J. E. Chambers, Q.-J. Meng, J. S. O’Neill, R. S. Edgar, and E. Derivery, Macromolecular condensation buffers intracellular water potential, Nature 623, 842 (2023).

[70] J. Bloustine, T. Virmani, G. M. Thurston, and S. Fraden, Light scattering and phase behavior of lysozyme-poly(ethylene glycol) mixtures, Phys. Rev. Lett. 96, 087803 (2006).

[71] Y. Osaka, R. Okamoto, T. Sumi, K. Koga, H. Imamura, T. Shirai, and Y. Isogai, Mechanism of polymer molecular weight-dependent suppression and promotion of liquid–liquid phase separation of a protein solution by the addition of polymer, The Journal of Chemical Physics 162, 051101 (2025), https://pubs.aip.org/aip/jcp/article-pdf/doi/10.1063/5.0245398/20375307/05110115.0245398.pdf.

[72] A. Kamo, A. Nikoubashman, and M. Yanagisawa, Impact of polydispersity on phase separation: Insights from polyethylene glycol and dextran mixtures, The Journal of Physical Chemistry B 129, 3263 (2025), pMID: 40080692, 10.1021/acs.jpcb.4c08640.

[73] V. A. Parsegian, R. P. Rand, and D. C. Rau, Osmotic stress, crowding, preferential hydration, and binding: A comparison of perspectives, Proceedings of the National Academy of Sciences 97, 3987 (2000), https://www.pnas.org/doi/pdf/10.1073/pnas.97.8.3987.

[74] R. D. A. Alvares, A. Hasabnis, R. S. Prosser, and P. M. Macdonald, Quantitative detection of pegylated biomacromolecules in biological fluids by nmr, Analytical Chemistry 88, 3730 (2016).

[75] M. I. Sanchez and A. Y. Ting, Directed evolution improves the catalytic efficiency of tev protease, Nature methods 17, 167 (2020).

[76] P. Zhou, A. A. Lugovskoy, and G. Wagner, A solubility-enhancement tag (set) for nmr studies of poorly behaving proteins, Journal of biomolecular NMR 20, 11 (2001).

